# Effects of age and noise on tympanal displacement in the Desert Locust

**DOI:** 10.1101/2023.09.14.557730

**Authors:** Thomas T Austin, Charlie Woodrow, Fernando Montealegre-Z, Ben Warren

## Abstract

Insect cuticle is an evolutionary-malleable exoskeleton that has specialised for various functions. Insects that detect the pressure component of sound bear specialised sound-capturing tympani evolved from cuticular thinning. Whilst the outer layer of insect cuticle is composed of non-living chitin, its mechanical properties change during development and aging. Here, we measured the displacements of the tympanum of the desert Locust, *Schistocerca gregaria*, to understand biomechanical changes as a function of age and noise-exposure. We found that the stiffness of the tympanum decreases within 12 hours of noise-exposure and increases as a function of age, independent of noise-exposure. Noise-induced changes were dynamic with an increased tympanum displacement to sound within 12 hours post noise-exposure. Within 24 hours, however, the tone-evoked displacement of the tympanum decreased below that of control Locusts. After 48 hours, the tone-evoked displacement of the tympanum was not significantly different to Locusts not exposed to noise. Tympanal displacements reduced predictably with age and repeatably noise-exposed Locusts (every three days) did not differ from their non-noise-exposed counterparts. Changes in the biomechanics of the tympanum may explain an age-dependent decrease in auditory detection in tympanal insects.

## Introduction

Animals that sensitively detect sound evolved specialised structures that capture sound energy. Some insect sound detectors capture the particle velocity component of sound with antennae, such as fruit flies and mosquitos, whilst others detect the pressure component of sound with specialised tympani, such as crickets and locusts. Vertebrate sound receivers are exclusively tympanal but even in invertebrates, tympani are the most common form of sound receiver (Yager, 1999). The tympanum of the desert Locust has two distinct regions of different thickness and stiffness which are straddled by Müller’s organ on its interior surface. The tympanum’s thin membrane is stiffer and detects high frequencies whereas the thick membrane is less stiff and detects low frequencies. Three groups of auditory neurons attach to the different parts of the tympanum to exploit differences in mechanical tuning. As such, Müller’s organ can detect frequencies from 200 Hz to at least 40 kHz (Römer, 1976). How does the mechanical function of the tympanum change as a function of age and noise exposure?

Longitudinal measurements of the human middle ear offer a good understanding of how sound receivers change as a function of age and noise-exposure. The mechanical properties of the middle ear change with age (Ruah et al., 1991) and could explain some age-related hearing loss (Corso, 1992, Rosenhall et al., 1990). Elderly humans with abnormal tympanometry are ∼50% more likely to have age-related hearing loss (Sogebi, 2015) and middle ear admittance (compliance) tends to decrease from the age of 20 to 40 (Wada et al., 1993). A consistent finding is no further changes in middle ear properties after the age of 40. This includes tympanal static admittance, typanometric peak pressure (the pressure at which there is greatest absorption of acoustic energy in the middle-ear), or resonance frequency of the middle ear (Uchida et al., 2000; Sinha et al., 2021) and acoustic conductance (Thompson et al., 1979).

Sound receivers of insects also change as a function of age and noise. For the sound receiver of the Locust, sound-evoked tympanal displacements decrease with age (Gordon and Windmill, 2015) which is concurrent with an age-related decrease in Müller’s organ function (Blockley et al., 2022). Directly after extended (24 hour) noise-exposure the Locust’s sound-evoked tympanal displacements dramatically increase (Warren et al., 2020) but recover to normal amplitudes 48 hours later for shorter (12 hour) noise-exposure. In contrast the passive stiffness of the antennal sound receiver of the fruit fly is robust to aging in spite of deterioration of auditory organ function (Keder et al., 2020). In response to noise-exposure changes in the antennal mechanics are determined by the active motility of the auditory neurons of Johnston’s organ (Boyd-Gibbins et al., 2021).

Electrophysiological performance and morphology of the Locust Müller’s organ is well characterised as a function of age (Blockley et al., 2022). The function of individual auditory neurons is well maintained but the auditory nerve response to sound decreases with age. In response to repeated noise-exposure (every three days) the function of the individual auditory neurons steadily deteriorates and the deterioration of auditory nerve function accelerates. For very old Locusts, age-related auditory decline dominates such that the difference between repeatably noise-exposed aged Locusts and control aged Locusts narrows. This echoes findings in humans, where hearing loss in repeatedly noise-exposed workers is most different to non-noise-exposed workers in middle-life (Corso, 1992).

Age and noise effects on the Locusts’ Müller’s organ (Blockley et a., 2022) could be explained by changes of the tympanum. Noise-exposure in the desert Locust has focused on Group-III auditory neurons – tuned to ∼3 kHz (Jacobs et al., 1999, Warren and Matherson, 2018) - that attach, through sclerites, to the foot of the styliform body (Fig. 1). Here, we examine the interaction of both noise-induced and age-related auditory decline by measuring tympanal displacements at the foot of the styliform body in response to 3 kHz tones.

**Figure 1:**
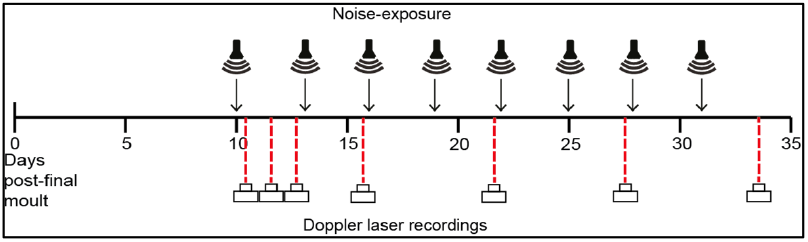
Timeline of noise-exposure and Doppler laser recordings.

## Methods

### Locust aging

Desert Locusts (*Schistocerca gregaria)* were raised in gregarious conditions in 60 cm^3^ cages, on a 12 hour light/dark cycle, at 36/25°C. Locusts were fed fresh wheat, grown in-house, and milled bran *ab libitum*. Ten days after their fifth and final moult, Locust’s wings were clipped to expose their abdominal tympani and Locusts entered the experimental pipeline where Locusts were housed in 20L aquarium tubs in crowded conditions and continued to be fed on fresh wheat and milled bran. Experimental day 0 corresponds with Locusts ten days post their last moult and experimental day 1 corresponds to directly after the Locusts’ first noise-exposure or mock exposure (control) to noise (Fig. 1).

### Noise-exposure and acoustic stimulation

Noise-exposed and control Locusts were placed in a cylindrical wire mesh cage (8 cm diameter, 11 cm height) placed directly under a speaker (Visaton FR 10 HM 4 OHM, RS Components Ltd). For the noise-exposed group, the speaker was driven by a function generator (Thurlby Thandar Instruments TG550, RS Components) and a sound amplifier (Monacor PA-702, Insight Direct) to produce a 3 kHz tone at 120 dB sound pressure level (SPL), measured at the top of the cage where Locusts tended to accumulate. Locusts were noise-exposed or mock noise-exposed (control) for 12 hours during the natural dark cycle every three days (day 0, 3, 6, 9, 12, 15, 18, 21). For example, Locusts recorded on day 13 had been noise-exposed once, those recorded on day 16 had been noise-exposed twice, day 22 had been noise-exposed four times (see Fig. 1).

For *in vivo* measurements of the tympanum, Locusts were mounted in natural dorso-ventral orientation following the removal of their hind legs and wings, and fixed to a copper platform using blue tac wrapped around their thorax and abdomen. A 3 kHz tone 50 ms duration with a 2ms rise time was produced with a waveform generator (SDG 1020, Siglent, China), and delivered via a stereo amplifier (SA1 power amplifier, Tucker-Davis Technologies, Alacchua, Florida) to a loudspeaker (MF1, Tucker-Davis Technologies, Alacchua, Florida) positioned 15 cm from the Locust. Sound amplitude was calibrated using a 1/8" microphone (Type 4138, Brüel & Kjaer, Germany) with built-in preamplifier (B&K 2670, Brüel & Kjær, Denmark), and a sound-level calibrator (Type 4237, Brüel & Kjaer, Denmark), via a conditioning amplifier (Nexus 2690-OS1, Brüel & Kjær, Denmark). Each Locust was stimulated with 50 3kHz tones at 30 -110 dB SPL and 55 ms duration (2 ms rise and fall times) in a randomised order.

### Biomechanical measurements of the tympanum with laser Doppler vibrometry

The Locust’s tympanum was orientated perpendicular to the micro-scanning Laser Doppler Vibrometer (PSV 500 with close-up unit and 150 mm lens, Polytec, Waldbronn, Germany). The inbuilt high-pass filter of the Polytec software was used to filter out slow movements, caused by breathing-related abdominal contractions. Displacement data of tympanum vibration was digitized using the PSV internal data acquisition board with a sampling rate of 512 kHz. Handling and measurements of each Locust took <5 min. Biomechanical measurements of the tympanum were carried out in an acoustic booth (IAC Acoustics, Series 120a, internal dimensions of 2.8 × 2.7 × 2 m) on a pneumatic vibration isolation table (Nexus Breadboard (B120150B), 1. 2 × 1.5 × 0.11 m, Thor Labs, USA). Displacements were calculated as the average peak-to-peak displacement.

### Data analysis

A power analysis was carried out to determine the number of Locusts required to achieve 95% power. The treatment of the Locust (noise-exposed or control) remained unknown to the experimenters, and all data remained blinded when analysing the data to avoid unconscious bias. Tympanal displacements were calculated by computing the peak-to-peak displacements for each sound amplitude for each Locust. In order to compare responses between the displacement of noise-exposed and control Locusts across sound pressure levels (SPLs), we adopted an approach first implemented in pharmacology research, which we have previously used to fit electrophysiological data (Warren et al., 2020). The “dose-response curves” represent SPL-auditory response curves. All models were fitted in R (Version 2.4.3), on a Windows PC running Windows 10. We fitted four-part log-linear models to the displacement of the tympanum data using the drm package in R when analysing displacement recordings (Ritz and Strebig, 2015), with displacement as the dependent variable with treatment (control or noise-exposed) and SPL as the independent variables. The t and p-values are reported for each model parameter: Hill coefficient (steepness of slope), maximal asymptote (maximum displacement), and inflection point (dB SPL at the steepest part of the slope) are shown on each graph in **Fig. 2**. The equation of the four-parameter log-linear fit is as follows:

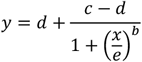

Where Y is displacement, b is the slope at the inflection point, c is the lower asymptote, d is the higher asymptote, e is the SPL (or X value) producing a response halfway between b and c. The drm package was also used to compute t and p-values when comparing four-part log-linear models. To test whether the factor of treatment (control or noise-exposed) significantly affected displacement, we compared the aforementioned model to a model in which treatment was omitted as an independent variable, using the anova function (Ritz et al., 2015). F statistics of the log-linear model fits were computed by excluding the treatment of interest (for example: control or noise-exposed) as an independent variable, and performing an anova between the models with and without this treatment. Higher F statistics donate a stronger effect of the treatment. Mixed effect linear models were created for each sound amplitude, introducing sex as a random factor, using the package *LME4* (Bates et al., 2015) in R Studio (version 1.4.1106).

**Figure 2:**
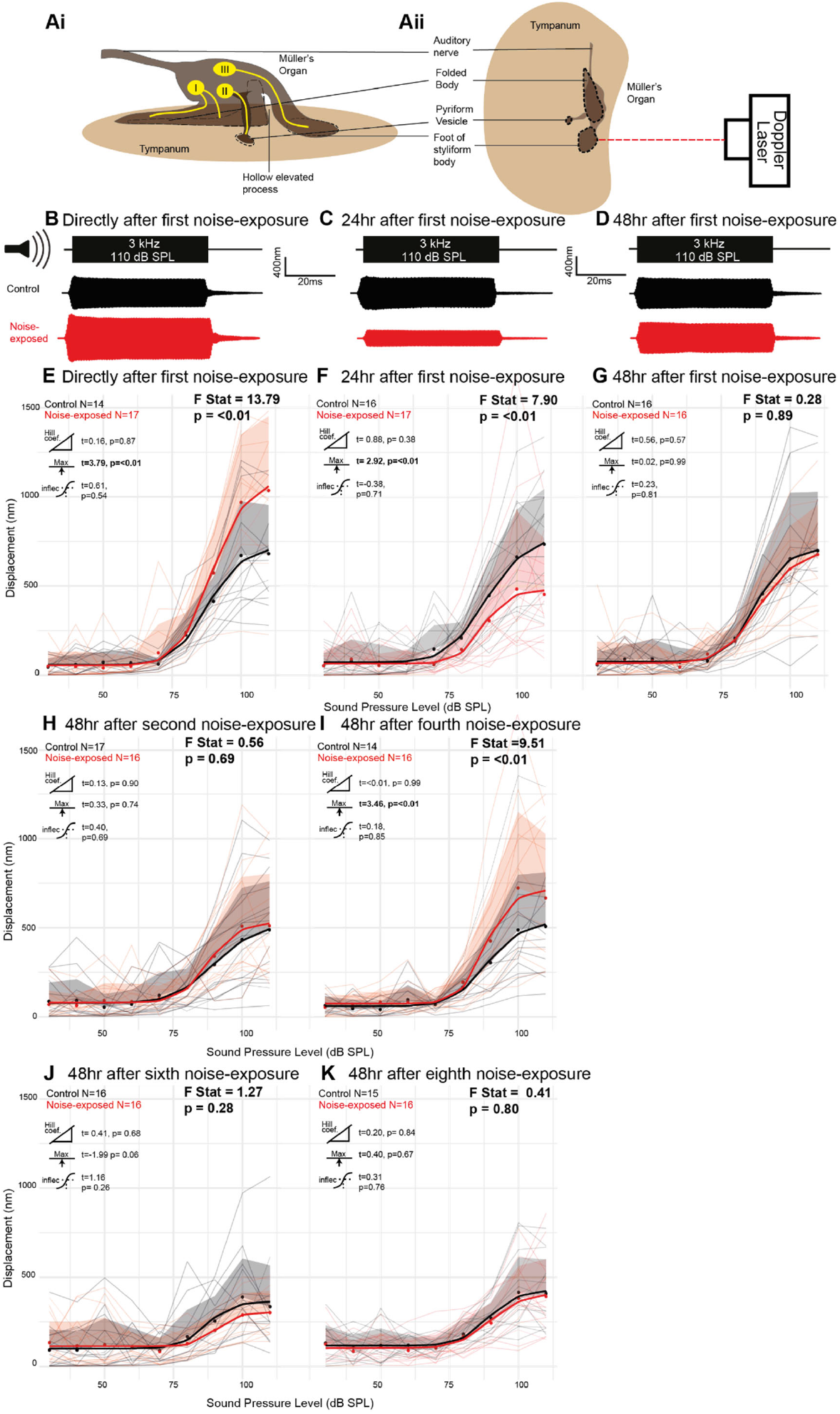
Doppler laser measurements of tone-evoked tympanal displacements from Locusts with and without noise-exposure. **Ai**. Schematic of Müller’s organ from internal oblique persepctive. **Aii** Schematic of recording position on the external surface of the tympanum. **B**. Displacement of the styliform body immediately after noise-exposure, **C**. 24 hours after noise-exposure, and **D**. 48 hours after mock noise exposure (control, black) and noise-exposure (red). **E**. Average displacements of the tympanum at the foot of the styliform body in response to 3 kHz pure tones for control (black trace) and noise-exposed (red trace) for Locusts recorded immediately after noise-exposure. The average tympanal displacements of individual Locusts are plotted as thin grey or thin red lines for control and noise-exposed groups. Dots are the average for each treatment at each SPL. The positive standard deviation is displayed in grey and red shaded areas. Log-linear fits are overlaid with thick black and red lines for control and noise-exposed. The extent of the difference between control and noise-exposed on each day is represented by the F-statistic, which compares the four-part log-linear models of control and noise-exposed cohorts. The hill coefficient, maximum asymptote and inflection point are compared between the two groups with a linear model. **F**. Control and noise-exposed Locust tone-evoked tympanal displacements 24 hours after noise-exposure **G**. 48 hours after noise-exposure. **H**. 48 hours after second noise-exposure. **I**. 48 hours after fourth noise-exposure. **J**. 48 hours after sixth noise-exposure. **K**. 48 hours after eighth noise-exposure.

## Results

Locusts were exposed to noise or mock noise-exposure every three nights (**Fig. 1**). Contactless laser Doppler vibrometry was used to measure *in vivo* tone-evoked tympanal displacements from the foot of the styliform body (where the sclerites of type III auditory neurons attach) in response to 3 kHz tones (**Fig. 2Aii**). In response to sound stimulation the styliform body on the tympanum vibrated at 3 kHz (**Fig. 2B**). Sound-evoked displacements of the tympanum increased directly after noise-exposure (**Fig. 2B**) but then decreased 24 hours later (**Fig. 2C**) before matching their age-matched controls 48 hours later (**Fig. 2D**). Tympanal displacements increased as a function of sound amplitude, with responses above the noise floor starting at ∼70 dB SPL and saturating above 100 dB SPL (**Fig. 2E-K**). To quantitively analyse responses of the tympanum we averaged tympanal displacements across sound amplitudes for noise-exposed and control Locusts and fitted the data with four-part log-linear models. Using this approach, we found that directly after noise-exposure maximal displacements are higher in noise-exposed Locusts (**Fig. 2E**: Fitted models F stat = 13.79, higher asymptote t_(543)_= 3.79). However, 24 hours after noise-exposure, tympanal displacements of the noise-exposed Locusts at sound amplitudes above 90 dB SPL shift to below that of control (**Fig. 2F:** Fitted models F stat = 7.90, higher asymptote t_(575)_= 2.92). After 48 hours, there was no significant difference in the tympanal displacements of the noise-exposed and control Locusts (**Fig. 2G**). Tympanal displacements of control groups not exposed to noise were similar directly after, 24 hours after and 48 hours after the first mock noise-exposure, (**Fig. 2E-G**, F stat = 0.304, p = 0.88; F stat = 0.140, p = 0.97; F stat = 0.173 p = 0.95).

We then examined tympanal displacement 48 hours after repeated noise-exposures in older Locusts (**Fig. 1)**. There was no difference between noise-exposed and control Locusts 48 hours after the second (**Fig. 2H**), sixth (**Fig. 2J**), and eighth (**Fig. 2K**) noise-exposure. This matches that recorded after the first noise-exposure (**Fig. 2G**). However, tympanal displacements of the noise-exposed Locusts tympanum were larger 48 hours after the fourth noise-exposure (**Fig. 2I**, F stat = 9.51, higher asymptote t_(507)_= 3.46).

Tympanal displacement, at higher sound amplitudes (90 dB SPL and 110 dB SPL) decreased significantly with age for both noise-exposed (**Fig. 3Di** 110 dB SPL: t_(74)_= -14.28, p =0.001; and **Fig. 3Dii** 90 dB SPL: t_(74)_= -8.78, p = 0.001) and control Locusts (**Fig. 3Ci** 110 dB SPL: t_(76)_= -13.53, p <0.001; and (**Fig. 3Cii** 90 dB SPL: t_(76)_= -7.22, p = 0.004). There was no change in displacement with age at 80 dB SPL (control: t_(76)_= -0.72, p = 0.63, and noise-exposed: t_(74)_= -1.37, p = 0.28), or lower sound amplitudes.

**Figure 3:**
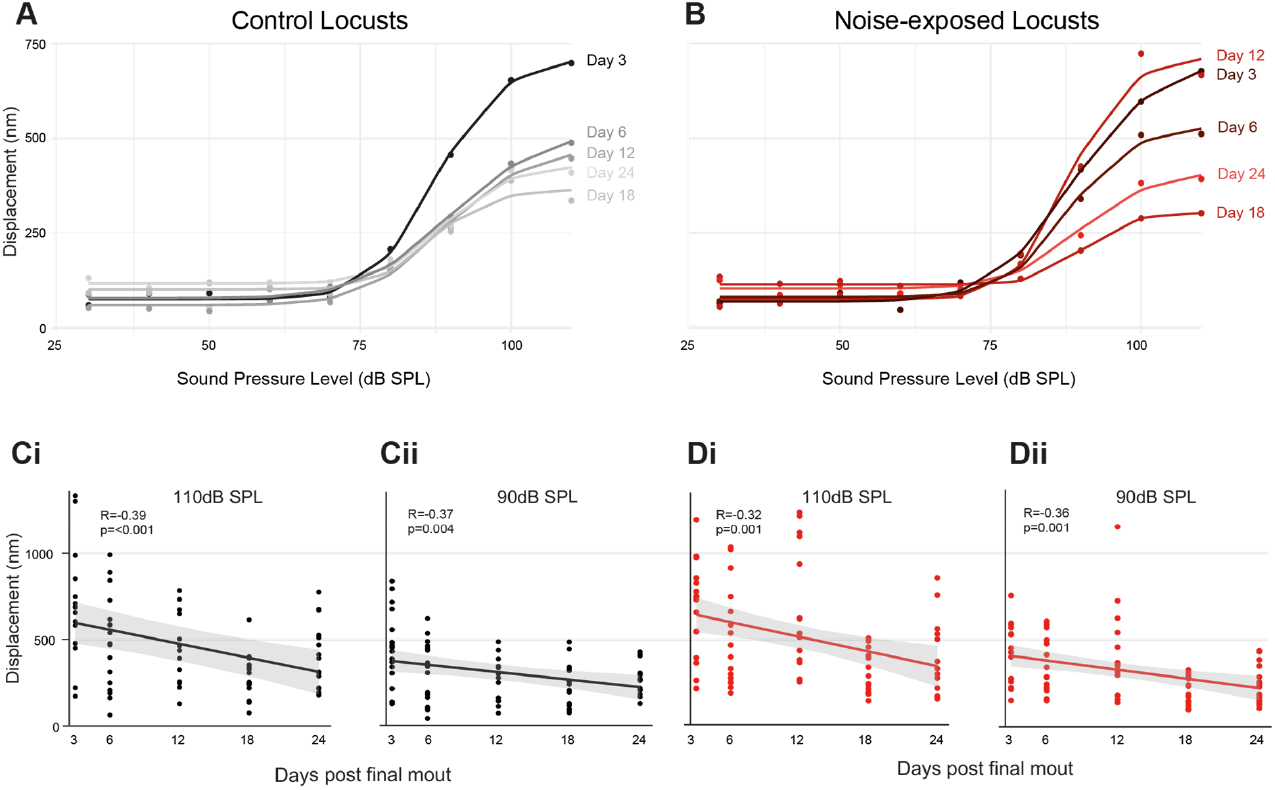
Doppler laser measurements of average tone-evoked tympanal displacements as a function of age for control and noise-exposed Locusts. **A**. Average tympanal displacement (dots) at each age for control Locusts fitted with log-linear functions (lines). **B**. Average tympanal displacement (dots) at each age for noise-exposed Locusts fitted with log-linear functions (lines). **C**. Tympanal displacement as a function of age for control locusts at **Ci** 110 dB SPL and **Cii** 90 dB SPL. Linear regression was overlaid with 95% confidence interval shaded. **D**. Tympanal displacement as a function of age for noise-exposed locusts at **Di** 110 dB SPL and **Dii** 90 dB SPL.

## Discussion

### Tympanal displacement decreases with age

Displacement of the tympanum at the styliform body decreases with age (**Fig. 3A**). This is similar to the decrease in displacement at the pyriform vesicle (Gordon and Windmill, 2015), between Locusts 13-days post final moult and older Locust groups ranging from 21-42-days - post final moult. Locusts do not moult again after their fifth and final moult but continue to add new layers of cuticle in a circadian and temperature-dependent cycle (Neville, 1963). This causes cuticle to harden in a process called sclerotization which impacts its mechanical properties (Anderson, 2010). Mechanical properties of Locust cuticle continue to change with age (Parle and Taylor, 2017), with the thickness and strength of the cuticle of the tibia increasing rapidly over the first 21 days post final moult. Sclerotization effects the function of other tissues of insects, such as in the tarsal pads in cockroaches, which harden and lose elasticity with age (Ridgel and Ritzmann, 2003, 2005, Zhou et al., 2015). Older animals are also less capable of restoring mechanical strength after insult but still deposit chitin and other cuticular proteins (O’Neill et al., 2019). In Müller’s Organ, vital genes involved in cuticle formation such as chitin and integrin have increased expression in younger Locusts (Austin et al., 2023), likely due to cuticular changes that accompany sclerotization after their final moult. Therefore, tympanal structural composition is likely to change with age, perhaps contributing to the decrease in compliance observed here and in Gordon and Windmill (2015). As suggested in Gordon and Windmill (2015) this could decrease the elasticity of the tympanal membrane, or the air sacs or the neuronal attachment points. In humans, although the tympanum histological and morphological structure deteriorates (Ruah et al., 1991; Etholm et al., 1974) its function is well maintained (Uchida et al., 2000; Sinha et al., 2021).

### Tympanal displacement increases with noise-exposure

Here, the displacement of the tympanum increases within hours after a 12-hour noise-exposure (**Fig. 2E**). After a 24-hour noise-exposure tympanal displacement also increases but across sound amplitudes from 50-100 dB SPL (Warren et al., 2020). The difference found here is limited to higher sound amplitudes, perhaps due to the 12-hour shorter noise-exposure. We discuss three possible mechanisms that causes changes in tympanal displacement after noise exposure.

Passive mechanical changes could explain a noise-induced increase in displacement. Noise-exposure could cause disruption of cytoskeletal structures attaching the tympanum to the cuticle and the tympanum to Müller’s organ, or damage to the cuticular structure of the tympanum itself. Genes involved in cuticle production and remodelling are differentially expressed in Locusts after noise-exposure (French and Warren, 2021), suggesting damage to the cuticle that is repaired.

The human ear has middle ear muscles that contract during high intensity sounds, that increase middle ear impedance, presumably to protect from acoustic overexposure (Mukerji et al., 2010). Anterior to the desert Locust’s tympanum is a spiracle with a muscle that controls the opening and closing at the spiracle entrance. The auditory nerve loops through the spiracle muscle and could alter stiffness of the tympanum by altering the tension of the connection of the auditory nerve to Müller’s organ. If the spiracle muscle is activated during acoustic stimulation, it could become overstimulated and reduce tympanal tension. The decrease in muscle tension could increase compliance of the tympanum. Muscle protein transcripts (troponin and myosin heavy chain) were reduced following noise-exposure (French and Warren, 2021), lending support to the hypothesis that a reduced muscle tension could increase displacements of the tympanum.

Insect ears react to low sound pressure levels nonlinearly (Göpfert and Robert, 2001), as found in other animals (Manley et al., 2001), including humans (Kemp, 1978). Nonlinear mechanical amplification helps to mechanically amplify quiet sounds and, in insects, is thought to be powered by dynein ATPases in the cilia of auditory neurons that interact with transduction channels (Albert et al., 2007; Warren et al., 2010). Clear evidence of active amplification is spontaneous movements of sound receivers above the thermal noise floor (Göpfert and Robert, 2001; Göpfert et al., 2005; Mhatre and Robert, 2013). Other manifestations of an active auditory system include echoes (otoacoustic emissions) of sounds played into an auditory system (Kemp, 1978) detectable my microphones. Although no such active movements have been found in the tympanum of the desert Locust, electrical stimulation of the auditory nerve resulted in reversible changes in (distortion product) otoacoustic emissions suggesting an active process (Möckel et al., 2007). Disruption of these mechanical properties immediately after noise-exposure could cause the increase in displacement found here.

### Tympanal displacement decreases in older Locusts regardless of noise-exposure

Although we find an initial increase in displacement immediately after noise-exposure, 24 hours later, tympanal displacement decreases significantly below the level of control animals (**Fig. 2F**). This decrease in displacement has also been found previously in Locusts 24 hours post-noise-exposure (Warren et al., 2020). This could be a homeostatic mechanism to recover to normal levels of displacement, however this mechanism overcompensates, resulting in a transient “overshoot”. This renders the tympanum of recovering noise-exposed Locusts temporarily less compliant than control Locusts. Locusts repair cuticular injuries with targeted deposition of cuticular material, regardless of age, at least in leg tibia (Parle et al., 2016, O’Neill et al., 2019). A similar mechanism is supported in the tympanum as glycoproteins involved in thickening of the endocuticle are upregulated in response to noise-exposure (French and Warren, 2021), and these may thicken and harden the tympanal membrane. Levels of displacement then return to the levels of the control Locust tympanum after 48 hours (**Fig. 2F**). Homeostatic maintenance of tympanal thickness and mass is likely key to preserve frequency-discriminating travelling waves in the Locust (Malkin et al., 2014), which would have large survival value.

### Tympanal displacements are typically normal 48 hours after noise-exposure, regardless of age

The displacement of the Locust’s tympanum 48 hours after noise-exposure is typically similar to control Locusts not exposed to noise, regardless of age. We assume that older Locusts undergo the same initial increase in tympanal displacement after noise-exposure (although we do not measure it) and then recover within the next 48 hours. In support of this age resilience, even older Locusts deposit chitin in a site-specific manner to repair and maintain cuticular integrity (O’Neill et al., 2019). Cuticle repair could be at the tympanum connection to the cuticular rim or more specific within the sclerites of Müller’s organ.

After a fourth noise-exposure, 22 days post-final moult, the Locusts tympanal displacement is higher than their age-matched controls (**Fig. 2I**). It is possible that the large individual variation between Locust tympanal displacements was due to a biological noise and individual variation, although this study was well-powered (at 95% power). It is also possible that the larger displacements of noise-exposed Locusts after the fourth noise-exposure reflect the large “middle-aged” difference found in the electrophysiological response of Müller’s organ (Blockley et al., 2022). This difference then narrows, possibly as age-related deterioration begins to dominate over noise-induced auditory loss.

